# Increased calcidiol level in redhaired people: Could redheadedness be an adaptation to temperate climate?

**DOI:** 10.1101/2019.12.30.890889

**Authors:** Jaroslav Flegr, Kateřina Sýkorová, Vojtěch Fiala, Jana Hlaváčová, Marie Bičíková, Ludmila Máčová, Šárka Kaňková

**Author notes:** **Corresponding author:** Jaroslav Flegr, Laboratory of Evolutionary Biology, Department of Philosophy and History of Sciences, Faculty of Science, Charles University, Vinicna 7, 128 00 Prague 2, Czech Republic.

## Abstract

About 1–2% of European population are redhaired, meaning they synthesize more pheomelanin than eumelanin, the main melanin pigment. Several mutations could be responsible for this phenotype. It has been suggested that corresponding mutations spread in Europe due to a founder effect shaped either by a relaxation of selection for dark, UV-protective phenotypes or by sexual selection in favor of rare phenotypes. In our study, we investigated the levels of vitamin D precursor calcidiol and folic acid in the blood serum of 73 redhaired and 130 non-redhaired individuals. In redhaired individuals, we found higher calcidiol concentrations and approximately the same folic acid concentrations as in non-redhaired subjects. Calcidiol concentrations correlated with the intensity of hair redness measured by two spectrophotometric methods and estimated by participants themselves and by independent observers. In non-redhaired individuals, calcidiol levels covaried with the amount of sun exposure and intensity of suntan while in redhaired individuals, this was not the case. It suggests that increased calcidiol levels in redhaired individuals are due to differences in physiology rather than in behavior. We also found that folic acid levels increased with age and the intensity of baldness and decreased with the frequency of visiting tanning salons. Our results suggest that the redhaired phenotype could be an evolutionary adaptation for sufficient photosynthesis of provitamin D in conditions of low intensity of UV-B radiation in central and northern parts of Europe.

## Introduction

On average, less than 2% of all Europeans (but 6–13% of population of Ireland, Wales, and Scotland) express the redhaired phenotype (Hooton, 1940; Sunderland & Barnicot, 1956). Mutations in the gene for receptor protein *MCIR* responsible for the absence or low levels of eumelanin in the affected subjects probably spread in human populations after the arrival of modern *Homo sapiens* to Europe. Nevertheless, the most common allele, Val92Met, seems to have introgressed into our gene pool from *Homo neanderthalensis* (Ding et al., 2014). It has been speculated that these alleles spread due to sexual selection, in particular by selection in favor of a rare phenotype (Frost, 2006; Frost, Kleisner, & Flegr, 2017). Many anecdotal observations (Chen et al., 2017; Liem, Hollensead, Joiner, & Sessler, 2006; Missmer et al., 2006; Somigliana et al., 2010; Tell-Marti et al., 2015) and one systematic largescale study (Frost et al., 2017) reveal that redhaired persons, especially women, tend to suffer from various symptoms of impaired health and from a higher frequency of certain diseases, including colorectal, cervical, uterine, and ovarian cancer than their non-redhaired peers. It has been suggested that the resulting selection against redhaired individuals counterbalances the positive sexual selection in favor of redhaired women, thereby maintaining the corresponding alleles at a low but stable frequency (Frost et al., 2017). Another study which used a similar population later showed that it is not the red hair as such but rather the pale skin frequently associated with redhaired phenotype that is responsible for the observed signs of impaired health of redhaired persons (Flegr & Sykorova, 2019). Pale skin can be the result of either congenitally low eumelanin concentrations in the skin or a sign of absence of suntan, usually due to avoidance of sun exposure (A. T. Slominski, Kim, Li, & Tuckey, 2016). The authors suggest that the impaired health (Skobowiat, Postlethwaite, & Slominski, 2017; A. Slominski & Postlethwaite, 2015) observed primarily in pale-skinned individuals and secondarily also in many redheaded persons is caused either by photolysis of folic acid in naturally pale individuals or by insufficient photosynthesis of vitamin D in persons who are pale due to avoidance of sun exposure. No direct data concerning the concentration of folic acid in pale-skinned or redhaired participants of that study (Flegr & Sykorova, 2019) were available. Rather surprisingly, we have not been able to find information about vitamin D and folic acid concentrations in redhaired individuals elsewhere in scientific literature either.

The aim of the present case-control study performed on a population of 203 subjects (73 of whom are redhaired) was to test the proposed hypotheses by searching for possible correlations between the intensity of natural hair redness, natural and by sun exposure acquired skin tone, and calcidiol and folic acid concentrations. In previous studies, the intensity of hair redness was rated by subjects themselves. To check the reliability of such data and their usefulness for future studies, we compared self-rated hair redness, redness rated by two independent observers, and exact measurements acquired by two different spectrophotometric methods.

## Materials and Methods

The project included a laboratory investigation which took place at the Faculty of Science of Charles University in Prague on September 17 – October 3, 2018. The second part, an online questionnaire survey, was completed by the same set of participants within the following 35 days.

### Participants

Participants were recruited mostly via a Facebook-based snowball method. Initially, an invitation to participate in a “study of health and personality of redheads” was posted on the timeline of the Facebook page Labbunnies, an approximately 18,000-strong group of Czech and Slovak nationals willing to participate in evolutionary psychology experiments. Further recruitment of redheads was carried out by invitations on Facebook, selective invitation of registered members of Labbunnies who completed our earlier questionnaires on the scale of redheadedness and scored four to six on a six-point scale, and by handing out flyers in the streets of Prague to people looked like natural redheads. We invited only people who confirmed that they had not dyed or bleached their hair for at least six months. This enabled us to measure natural hair color near the hair roots. Only subjects who provided informed consent were included in the study. In the end, we assembled a sample of 110 women and 93 men. Participants received no remuneration, only a commemorative badge and a haircare gift set (costing 53 CZK, that is app. 2.3 USD). The project was approved by the Ethics Committee of the Faculty of Science, Charles University (No. 2018/30).

### Experimental design

Participants were instructed to wash their hair the evening before or morning of the day of the laboratory measurement and to refrain from using any post-shampoo products. At the beginning of the session, participants obtained a paper questionnaire, which they could complete while waiting for individual measurements. First, participants were tested with an electronic dynamometer (not part of the present study). Then we measured the natural red color of participants’ hair and their skin hue by using a spectrophotometer to obtain a standardized scale of redheadedness and skin hue. Subsequently, participants were tested with a mechanical dynamometer. While the dynamometer and spectrophotometer measurements were performed in two separate rooms, two observers (a woman and a man) independently rated the intensity of subjects’ redheadedness and freckledness using an ordinal scale of zero to five. At the end of the laboratory part of the study, we asked participants if they consent to having a blood sample taken to determine the concentration of calcidiol and folic acid. The sampling was performed in an adjacent room by a qualified nurse. Several days after the laboratory part of the study, we sent all participants a link to another electronic questionnaire with a request to complete it within the following 35 days. After two rounds of e-mail reminders, 198 (97.5%) of participants completed this questionnaire.

### Questionnaires

All participants were asked to complete one printed questionnaire and one electronic questionnaire, distributed via Qualtrics platform, which aimed at collecting their basic anamnestic information and information related to their and their relatives’ hair and body pigmentation, as well as their tanning or sun-avoidance behaviors. Specifically, we asked the participants to rate the following:

– natural redness of their hair and hair color in childhood on a six-point scale anchored with “absolutely non-red” (code 1) – “bright red” (code 6);
– natural lightness of their hair and complexion on a six-point scale anchored with “very light” (code 1) – “very dark” (code 6);
– hair length on a four-point scale anchored with “very short, does not cover the forehead, ears, or neck” (code 1) – “medium length or long, covering forehead, ears, and neck, mostly worn loose” (code 4);
– intensity of baldness on a seven-point scale, where degrees were shown by black and white pictures (no responder chose code 7, the highest degree of baldness);
– current intensity of suntan on a six-point scale anchored with “no suntan” (code 1) – “very dark suntan” (code 6);
– tendency to tan to brown and tendency to tan to red on six-point scales anchored with “definitely not” (code 1) – “definitely yes” (code 6);
– intensity of chemical self-protection from sun by creams or oils with UV filters, intensity of self-protection by mechanical means (by shelters and clothing) on six-point scales anchored with “not at all” (code 1) – “yes, very carefully” (code 6);
– frequency of sun exposure on a seven-point scale anchored with “almost never” (code 1) – “over three hours a day” (code 7); no responder chose code 7;
– frequency of visits to tanning salons on a five-point scale anchored with “never” (code 1) – “yes, almost throughout the year” (code 5);
– frequency of taking vitamin D supplements on a six-point scale anchored with “never” (code 2) – “yes, almost constantly” (code 6). Here, responders could also check “I do not know” (code 1: “missing value”).

Participants were also asked whether they had red hair on other parts of their body (e.g. facial hair, body hair) and whether they had redhaired relatives (binary variables). The other two binary variables of red hair (no/yes) and red hair in childhood (no/yes) were obtained by splitting the corresponding ordinal variables (0: 1, 2 vs. 1: 3, 4, 5, 6). We also monitored potential confounding variables such as sex, age, and size of place of residence (six categories: <1000 inhabitants,1,000–5,000, 5,000–50,000, 50,000–100,000, 100,000–500,000, Prague or Bratislava).

### Measuring skin and hair pigmentation with a spectrophotometer

Measurements of the natural red color of participant’s hair and darkness or lightness of their skin tone was performed with a spectrophotometer (Ocean Optics FLAME-S). The device was white-calibrated using a WS-1 Diffuser Reflectance Standard. Before commencing the measurement, the experimenter asked if participant’s hair had been dyed and then cleaned the participant’s cheeks and forehead with a make-up removal pad. Then he took three spectrophotometric measurements of skin color on the inner upper arm of the less dominant hand (depending on participants’ self-reported handedness), one on the left cheek, one on the right cheek, and one on the forehead above nasal root. To measure hair color, the experimenter moved aside the crown hair to get to the hair in the occipital region and made sure that the scalp was not visible. Then he took three spectrophotometric measurements of hair color in different areas around the occipital region. The occipital region and inner upper arm are the areas least exposed to sunlight, which is why hair and skin color found there correspond most closely to the natural color. We used two methods to determine the total level of redheadedness. The first was Reed’s function (Reed, 1952):

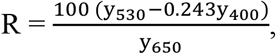

where y_400_, y_530_, and y_650_ are the arithmetical means of three measurements of percentage reflectance values at wavelengths in the subscript. The second was the CIE L*a*b* color space: it provided the a* parameter which ranges from −100 (green) to +100 (red) (Lozano, Saunier, Panhard, & Loussouarn, 2017; Vaughn, van Oorschot, & Baindur-Hudson, 2008).

### Measurements of calcidiol and folic acid concentrations

Calcidiol concentration was measured using High Performance Liquid Chromatography by ClinRep® Complete Kit for 25-OH-Vitamin D2/D3 (RECIPE Chemicals + Instruments GmbH, Munich, Germany). Folic acid was measured with ID-Vit®Folic acid microtiter plate kit (Immundiagnostik AG, Bensheim, Germany). After incubation at 37°C for 48 hours, the growth of *Lactobacillus rhamnosusis* was measured turbidimetrically at 620 nm using ELISA-reader Spark™ 10M (Tecan, Männedorf, Switzerland).

### Statistics

Statistica v. 10.0 was used to explore the data and R v. 3.3.1 (R Core Team, 2018) for confirmatory statistical tests. Associations of sex with age, and calcidiol and folic acid concentrations were estimated by a t-test and correlation of sex, age, and urbanization with all focal variables by a Kendall correlation test. Partial Kendall correlation test (R package ppcor 1.1 (Kim, 2015)) with age, urbanization, and in some analyses also sex as potential covariates was used for the main analysis. This multivariate nonparametric test allows for measuring the significance and strength of correlations between any combination of binary, ordinal, and continuous variables while controlling for any number of confounding variables. In the confirmatory part of the study, i.e. to test the hypothesized effect of redheadedness on calcidiol and folic acid concentrations, we performed a correction for multiple tests by Benjamini-Hochberg procedure with false discovery rate preset to 0.20 (Benjamini & Hochberg, 1995). In the exploratory parts of the study, we performed no correction for multiple tests.

## Results

The final population consisted of 110 women (mean age 27.4, SD 7.5) and 93 men (mean age 34.0, SD 9.0). The age difference between men and women was highly significant (t_180_=−3.92, p=0.0001, Cohen’s d = 0.563). Table 1 shows the descriptive statistics for our ordinal and binary data. Kendall correlation test showed that men and women, the old and the young, people residing in small and large settlements, and redhaired versus non-redhaired people differed in their responses to hair and body pigmentation-related variables as well as in behavioral variables related to sun exposure (Table 1, the last four columns). Average calcidiol concentrations were higher in 105 women (75.1 nmol/L, SD 22.8) than in 88 men (70.2 nmol/L, SD 21.8) but the difference was not statistically significant (t_186_ = 1.54, p = 0.124, Cohen’s d = −0.224). Folic acid concentrations were also non-significantly higher in 99 women (7.48 μg/L, SD 5.71) than in 78 men (7.11 μg/L, SD 4.78) (t_174_ = 1.54, p = 0.643, Cohen’s d = −0.069). Table 2 shows correlations between calcidiol and folic acid concentrations and age and urbanization. Except for a strong positive correlation between folic acid concentration and age (Tau = 0.210, p < 0.00001), none of these correlations reached the formal level of statistical significance.

**Table 1.** Distributions of responses of participants (or observers) to particular questions.

|  |  | 1 |  | 2 |  | 3 |  | 4 |  | 5 |  | 6 |  |  | Sex | Age | Urban. | Redness |
| --- | --- | --- | --- | --- | --- | --- | --- | --- | --- | --- | --- | --- | --- | --- | --- | --- | --- | --- |
|  |  | n | % | n | % | n | % | n | % | n | % | n | % |  |  |  |  |  |
| Urbanization | women | 11 | 10.00 | 14 | 12.73 | 9 | 8.18 | 2 | 1.82 | 1 | 0.91 | 73 | 66.36 | Tau | 0.072 | 0.082 | NA | 0.085 |
|  | men | 3 | 3.23 | 10 | 10.75 | 10 | 10.75 | 1 | 1.08 | 3 | 3.23 | 66 | 70.97 | P | 0.13 | 0.08 | NA | 0.07 |
| Hair redness | women | 50 | 45.45 | 16 | 14.55 | 7 | 6.36 | 17 | 15.45 | 13 | 11.82 | 7 | 6.36 | Tau | -0.088 | -0.082 | 0.085 | NA |
|  | men | 49 | 52.69 | 15 | 16.13 | 9 | 9.68 | 8 | 8.60 | 7 | 7.53 | 5 | 5.38 | P | 0.06 | 0.08 | 0.07 | NA |
| Redness childhood | women | 53 | 49.07 | 9 | 8.33 | 4 | 3.70 | 7 | 6.48 | 10 | 9.26 | 25 | 23.15 | Tau | -0.117 | -0.084 | 0.095 | 0.801 |
|  | men | 56 | 60.22 | 4 | 4.30 | 7 | 7.53 | 6 | 6.45 | 8 | 8.60 | 12 | 12.90 | P | **0.01** | 0.08 | **0.05** | **0.00** |
| Body hair redness | women | 70 | 63.64 | 40 | 36.36 |  |  |  |  |  |  |  |  | Tau | 0.089 | 0.053 | 0.100 | 0.680 |
|  | men | 51 | 54.84 | 42 | 45.16 |  |  |  |  |  |  |  |  | P | 0.06 | 0.26 | **0.03** | **0.00** |
| Red hair relatives | women | 71 | 65.74 | 37 | 34.26 |  |  |  |  |  |  |  |  | Tau | 0.013 | -0.016 | 0.062 | 0.523 |
|  | men | 60 | 64.52 | 33 | 35.48 |  |  |  |  |  |  |  |  | P | 0.79 | 0.73 | 0.19 | **0.00** |
| Hair redness observer 1 | women | 52 | 47.27 | 16 | 14.55 | 2 | 1.82 | 3 | 2.73 | 15 | 13.64 | 22 | 20.00 | Tau | -0.151 | -0.127 | 0.047 | 0.745 |
|  | men | 59 | 63.44 | 11 | 11.83 | 5 | 5.38 | 1 | 1.08 | 3 | 3.23 | 14 | 15.05 | P | **0.00** | **0.01** | 0.32 | **0.00** |
| Hair redness observer 2 | women | 5 | 4.55 | 12 | 10.91 | 18 | 16.36 | 33 | 30.00 | 24 | 21.82 | 18 | 16.36 | Tau | -0.103 | -0.087 | 0.010 | 0.574 |
|  | men | 11 | 11.83 | 15 | 16.13 | 14 | 15.05 | 22 | 23.66 | 18 | 19.35 | 13 | 13.98 | P | **0.03** | 0.07 | 0.83 | **0.00** |
| Hair darkness | women | 1 | 0.91 | 17 | 15.45 | 33 | 30.00 | 33 | 30.00 | 24 | 21.82 | 2 | 1.82 | Tau | 0.108 | 0.073 | -0.021 | -0.277 |
|  | men | 0 | 0.00 | 9 | 9.68 | 26 | 27.96 | 28 | 30.11 | 27 | 29.03 | 3 | 3.23 | P | **0.02** | 0.12 | 0.66 | **0.00** |
| Baldness | women |  |  |  |  |  |  |  |  |  |  |  |  | Tau | NA | 0.229 | 0.146 | -0.104 |
|  | men | 41 | 44.09 | 40 | 43.01 | 6 | 6.45 | 2 | 2.15 | 0 | 0.00 | 0 | 0.00 | P | NA | **0.00** | **0.04** | 0.15 |
| Hair length | women | 0 | 0.00 | 6 | 5.45 | 56 | 50.91 | 48 | 43.64 |  |  |  |  | Tau | -0.666 | -0.105 | -0.006 | 0.155 |
|  | men | 39 | 42.39 | 38 | 41.30 | 9 | 9.78 | 6 | 6.52 |  |  |  |  | P | **0.00** | **0.03** | 0.89 | **0.00** |
| Natural skin darkness | women | 22 | 20.00 | 49 | 44.55 | 26 | 23.64 | 12 | 10.91 | 1 | 0.91 | 0 | 0.00 | Tau | 0.161 | 0.123 | -0.098 | -0.406 |
|  | men | 13 | 13.98 | 31 | 33.33 | 29 | 31.18 | 16 | 17.20 | 3 | 3.23 | 1 | 1.08 | P | **0.00** | **0.01** | **0.04** | **0.00** |
| Suntan | women | 11 | 10.00 | 35 | 31.82 | 30 | 27.27 | 25 | 22.73 | 8 | 7.27 | 1 | 0.91 | Tau | 0.057 | 0.059 | -0.113 | -0.157 |
|  | men | 8 | 8.60 | 31 | 33.33 | 16 | 17.20 | 26 | 27.96 | 10 | 10.75 | 2 | 2.15 | P | 0.23 | 0.21 | **0.02** | **0.04** |
| Brown tanning | women | 21 | 19.09 | 25 | 22.73 | 18 | 16.36 | 18 | 16.36 | 21 | 19.09 | 7 | 6.36 | Tau | 0.083 | 0.077 | -0.036 | -0.407 |
|  | men | 12 | 12.90 | 20 | 21.51 | 16 | 17.20 | 15 | 16.13 | 22 | 23.66 | 8 | 8.60 | P | 0.08 | 0.10 | 0.44 | **0.00** |
| Red tanning | women | 18 | 16.36 | 24 | 21.82 | 15 | 13.64 | 13 | 11.82 | 22 | 20.00 | 18 | 16.36 | Tau | 0.001 | -0.016 | 0.150 | 0.432 |
|  | men | 10 | 10.75 | 22 | 23.66 | 18 | 19.35 | 14 | 15.05 | 17 | 18.28 | 12 | 12.90 | P | 0.99 | 0.73 | **0.00** | **0.00** |
| Freckledness observer 1 | women | 54 | 49.09 | 17 | 15.45 | 17 | 15.45 | 7 | 6.36 | 4 | 3.64 | 11 | 10.00 | Tau | -0.196 | -0.045 | 0.053 | 0.518 |
|  | men | 61 | 65.59 | 19 | 20.43 | 5 | 5.38 | 6 | 6.45 | 0 | 0.00 | 2 | 2.15 | P | **0.00** | 0.34 | 0.26 | **0.00** |
| Freckledness observer 2 | women | 5 | 4.55 | 12 | 10.91 | 18 | 16.36 | 33 | 30.00 | 24 | 21.82 | 18 | 16.36 | Tau | -0.150 | -0.023 | 0.082 | 0.542 |
|  | men | 11 | 11.83 | 15 | 16.13 | 14 | 15.05 | 22 | 23.66 | 18 | 19.35 | 13 | 13.98 | P | **0.00** | 0.63 | 0.08 | **0.00** |
| Protection by sun creams | women | 9 | 8.18 | 13 | 11.82 | 11 | 10.00 | 28 | 25.45 | 26 | 23.64 | 23 | 20.91 | Tau | -0.163 | -0.122 | 0.171 | 0.250 |
|  | men | 15 | 16.13 | 15 | 16.13 | 14 | 15.05 | 20 | 21.51 | 18 | 19.35 | 11 | 11.83 | P | **0.00** | **0.01** | **0.00** | **0.00** |
| Protection by shelters | women | 21 | 19.09 | 24 | 21.82 | 20 | 18.18 | 22 | 20.00 | 15 | 13.64 | 8 | 7.27 | Tau | -0.099 | -0.006 | 0.015 | 0.177 |
|  | men | 20 | 21.51 | 26 | 27.96 | 17 | 18.28 | 20 | 21.51 | 10 | 10.75 | 0 | 0.00 | P | **0.04** | 0.91 | 0.76 | **0.00** |
| Sun exposure | women | 3 | 2.73 | 1 | 0.91 | 13 | 11.82 | 17 | 15.45 | 27 | 24.55 | 49 | 44.55 | Tau | -0.020 | -0.019 | -0.061 | 0.001 |
|  | men | 0 | 0.00 | 1 | 1.08 | 6 | 6.45 | 17 | 18.28 | 37 | 39.78 | 32 | 34.41 | P | 0.67 | 0.69 | 0.20 | 0.99 |
| Tanning salons | women | 103 | 93.64 | 4 | 3.64 | 2 | 1.82 | 1 | 0.91 | 0 | 0.00 |  |  | Tau | -0.173 | -0.001 | 0.072 | -0.010 |
|  | men | 93 | 100.00 | 0 | 0.00 | 0 | 0.00 | 0 | 0.00 | 0 | 0.00 |  |  | P | **0.00** | 0.98 | 0.13 | 0.84 |
| Vitamin D supplements | women | 21 | 19.27 | 56 | 51.38 | 24 | 22.02 | 0 | 0.00 | 7 | 6.42 | 1 | 0.92 | Tau | -0.096 | 0.130 | 0.015 | -0.088 |
|  | men | 27 | 29.03 | 44 | 47.31 | 16 | 17.20 | 0 | 0.00 | 2 | 2.15 | 4 | 4.30 | P | **0.04** | **0.01** | 0.75 | 0.06 |

This table shows the distribution of responses of our subjects (and observers) to particular questions. The last four columns show the strength and significance (Tau and p) of Kendall correlations between variables listed in the first column and sex, age, urbanization, and intensity of hair redness (controlled for age and urbanization), respectively. Positive Tau means that men, older subjects, residents of larger cities, and more redhaired subjects provided higher codes of responses than women, younger subjects, residents of smaller cities, and less redhaired subjects (see Material and
methods). Significant correlations are printed in bold. Statistical significance below 0.005 is coded 0.00.

**Table 2.** The effects of variables related to body pigmentation and sun exposure behaviors on calcidiol and folic acid concentrations.

|  | ALL |  |  |  | MEN |  |  |  | WOMEN |  |  |  |
| --- | --- | --- | --- | --- | --- | --- | --- | --- | --- | --- | --- | --- |
|  | calcidiol |  | folic acid |  | calcidiol |  | folic acid |  | calcidiol |  | folic acid |  |
|  | Tau | p | Tau | p | Tau | p | Tau | p | Tau | p | Tau | p |
| Age | -0.061 | 0.210 | 0.210 | **0.000** | -0.070 | 0.339 | 0.113 | 0.148 | -0.015 | 0.820 | 0.285 | **0.000** |
| Urbanization | -0.062 | 0.201 | 0.018 | 0.722 | 0.024 | 0.739 | 0.028 | 0.723 | -0.120 | 0.072 | 0.016 | 0.819 |
| Hair redness | 0.142 | **0.004** | 0.002 | 0.968 | 0.138 | 0.060 | -0.134 | 0.086 | 0.149 | **0.027** | 0.089 | 0.195 |
| Hair redness binary | 0.229 | **0.000** | -0.017 | 0.737 | 0.222 | **0.002** | -0.172 | **0.028** | 0.239 | **0.000** | 0.077 | 0.265 |
| Redness childhood | 0.141 | **0.004** | 0.004 | 0.931 | 0.198 | **0.007** | -0.126 | 0.107 | 0.110 | 0.105 | 0.074 | 0.286 |
| Redness childhood binary | 0.173 | **0.000** | -0.010 | 0.851 | 0.252 | **0.001** | -0.167 | **0.033** | 0.124 | 0.064 | 0.099 | 0.151 |
| Body hair redness | 0.203 | **0.000** | -0.018 | 0.722 | 0.186 | **0.011** | -0.101 | 0.196 | 0.236 | **0.000** | 0.043 | 0.537 |
| Red hair relatives | 0.072 | 0.145 | -0.091 | 0.075 | 0.116 | 0.114 | -0.227 | **0.004** | 0.036 | 0.595 | 0.011 | 0.875 |
| Hair redness observer 1 | 0.192 | **0.000** | 0.007 | 0.884 | 0.165 | **0.025** | -0.112 | 0.154 | 0.206 | **0.002** | 0.062 | 0.369 |
| Hair redness observer 2 | 0.167 | **0.001** | -0.015 | 0.763 | 0.185 | **0.012** | -0.097 | 0.214 | 0.160 | **0.017** | 0.047 | 0.492 |
| Reflectance 400 | 0.128 | **0.009** | -0.011 | 0.833 | 0.159 | **0.030** | -0.118 | 0.131 | 0.114 | 0.090 | 0.084 | 0.225 |
| Reflectance 530 | 0.177 | **0.000** | 0.016 | 0.750 | 0.226 | **0.002** | -0.085 | 0.275 | 0.146 | **0.029** | 0.110 | 0.112 |
| Reflectance 650 | 0.204 | **0.000** | 0.015 | 0.768 | 0.231 | **0.002** | -0.062 | 0.429 | 0.179 | **0.008** | 0.082 | 0.235 |
| Hair redness R | -0.202 | **0.000** | -0.004 | 0.937 | -0.189 | **0.010** | -0.002 | 0.978 | -0.185 | **0.006** | 0.014 | 0.839 |
| Hair redness a* | 0.205 | **0.000** | 0.012 | 0.807 | 0.200 | **0.006** | -0.039 | 0.614 | 0.189 | **0.005** | 0.041 | 0.552 |
| Hair darkness | -0.117 | **0.016** | -0.006 | 0.899 | -0.132 | 0.071 | 0.110 | 0.159 | -0.097 | 0.147 | -0.091 | 0.185 |
| Hair length | 0.045 | 0.357 | 0.039 | 0.441 | -0.074 | 0.317 | -0.108 | 0.171 | 0.072 | 0.283 | 0.149 | **0.031** |
| Baldness | 0.048 | 0.511 | 0.208 | **0.008** | 0.048 | 0.511 | 0.208 | **0.008** | NA | NA | NA | NA |
| Natural skin darkness | -0.036 | 0.464 | 0.072 | 0.159 | 0.031 | 0.674 | 0.152 | 0.052 | -0.102 | 0.127 | 0.010 | 0.880 |
| Suntan | 0.151 | **0.002** | 0.067 | 0.189 | 0.191 | **0.009** | 0.079 | 0.311 | 0.111 | 0.099 | 0.074 | 0.286 |
| Brown tanning | -0.035 | 0.474 | 0.005 | 0.925 | -0.031 | 0.672 | 0.102 | 0.193 | -0.051 | 0.445 | -0.047 | 0.499 |
| Red tanning | 0.049 | 0.317 | 0.007 | 0.898 | 0.101 | 0.167 | -0.076 | 0.334 | 0.024 | 0.726 | 0.049 | 0.477 |
| Freckledness observer 1 | 0.120 | **0.014** | -0.024 | 0.631 | 0.062 | 0.395 | -0.154 | **0.049** | 0.142 | **0.034** | 0.033 | 0.628 |
| Freckledness observer 2 | 0.121 | **0.013** | -0.031 | 0.541 | 0.121 | 0.099 | -0.125 | 0.110 | 0.121 | 0.072 | 0.018 | 0.789 |
| Facial skin fairness | -0.066 | 0.173 | -0.016 | 0.746 | -0.202 | **0.006** | -0.136 | 0.082 | -0.087 | 0.194 | 0.031 | 0.658 |
| Arm skin fairness | -0.079 | 0.106 | 0.038 | 0.460 | -0.124 | 0.091 | -0.031 | 0.687 | -0.066 | 0.323 | 0.055 | 0.423 |
| Protection by sun creams | 0.043 | 0.373 | -0.032 | 0.534 | 0.087 | 0.238 | -0.144 | 0.066 | -0.009 | 0.898 | 0.023 | 0.734 |
| Protection by shelters | -0.079 | 0.107 | -0.039 | 0.439 | -0.084 | 0.253 | -0.107 | 0.173 | -0.086 | 0.200 | -0.006 | 0.936 |
| Sunbathing | 0.114 | **0.020** | 0.033 | 0.521 | 0.208 | **0.005** | 0.048 | 0.543 | 0.052 | 0.435 | 0.030 | 0.666 |
| Tanning salons | -0.016 | 0.743 | -0.140 | **0.006** | NA | NA | NA | NA | -0.043 | 0.526 | -0.209 | **0.002** |
| Vitamin D supplements | 0.003 | 0.956 | 0.026 | 0.659 | 0.010 | 0.913 | 0.065 | 0.493 | -0.007 | 0.928 | -0.010 | 0.894 |

*This table shows the strength and direction of partial Kendall correlations (controlled for age and urbanization) between variables listed in the first column (see Material and methods) and calcidiol and folic acid concentrations. Significant correlations are printed in bold.*

The effect of hair redness and other variables related to hair and body pigmentation as well as sun exposure behaviors on the concentration of calcidiol and folic acid in the serum was analyzed primarily with nonparametric partial Kendall correlations controlled for age and urbanization. Nevertheless, similar results were obtained also when sex, hair and skin tone (light to dark), and even the frequency of sun exposure and intensity of suntan were controlled for. Our results suggest that hair redness has the strongest effect on calcidiol concentrations (significant after correction for multiple tests) and a negligible effect on folic acid concentrations (see Figures 1–2, Table 2). The strongest correlation was observed when analyses used the binary variable hair redness obtained from hair redness estimated by the subjects on an ordinal scale of 1–6 split to 0 (responses 1 and 2) and 1 (responses 3–6). Nonetheless, effects of a similar strength were detected when the intensity of hair redness was measured spectrophotometrically and that held regardless of which index, including raw reflectance at 650 nm, of hair redness was applied.

**Fig. 1.**
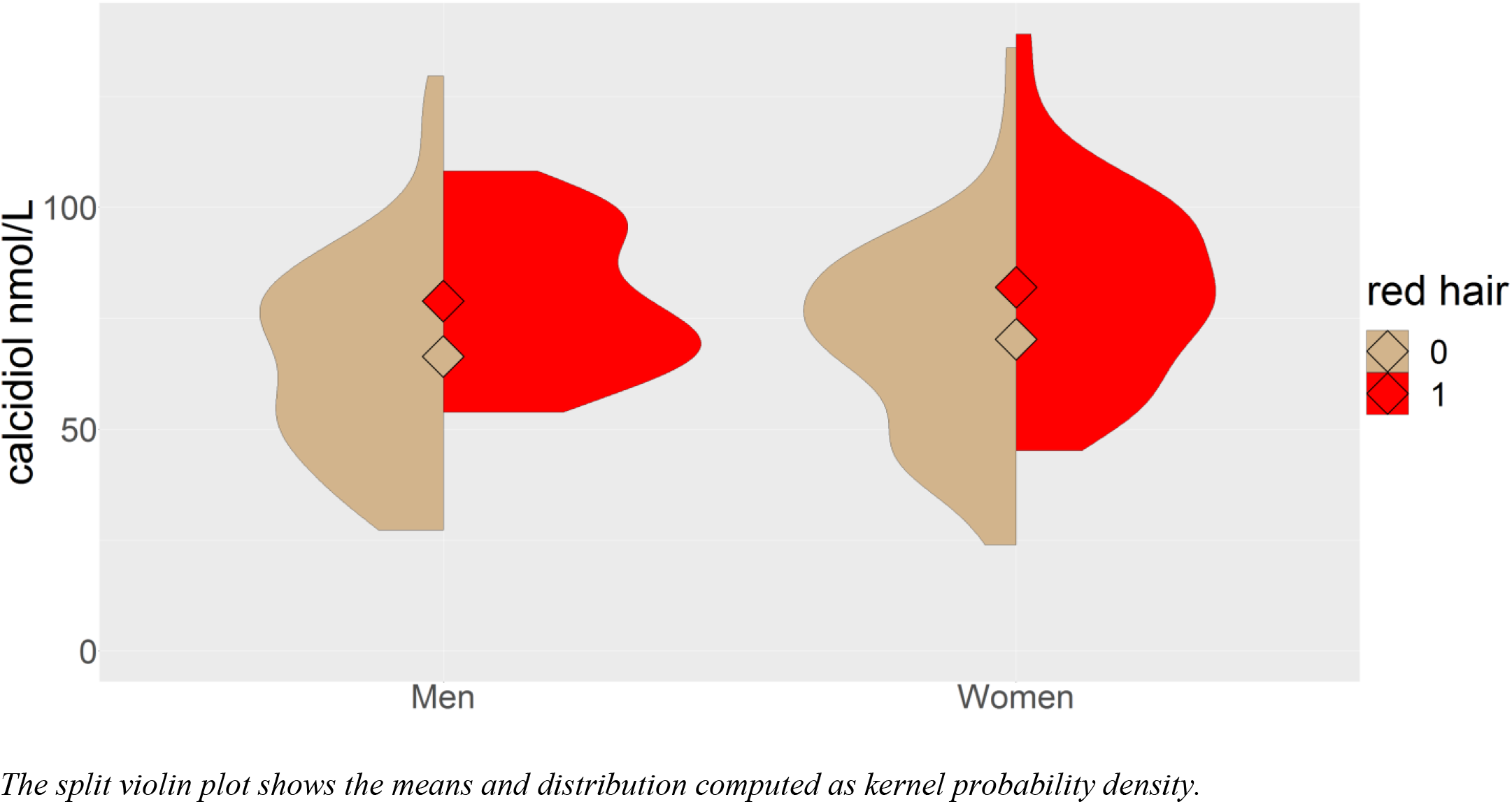
The effect of sex and red hair color on calcidiol concentration.

**Fig. 2.**
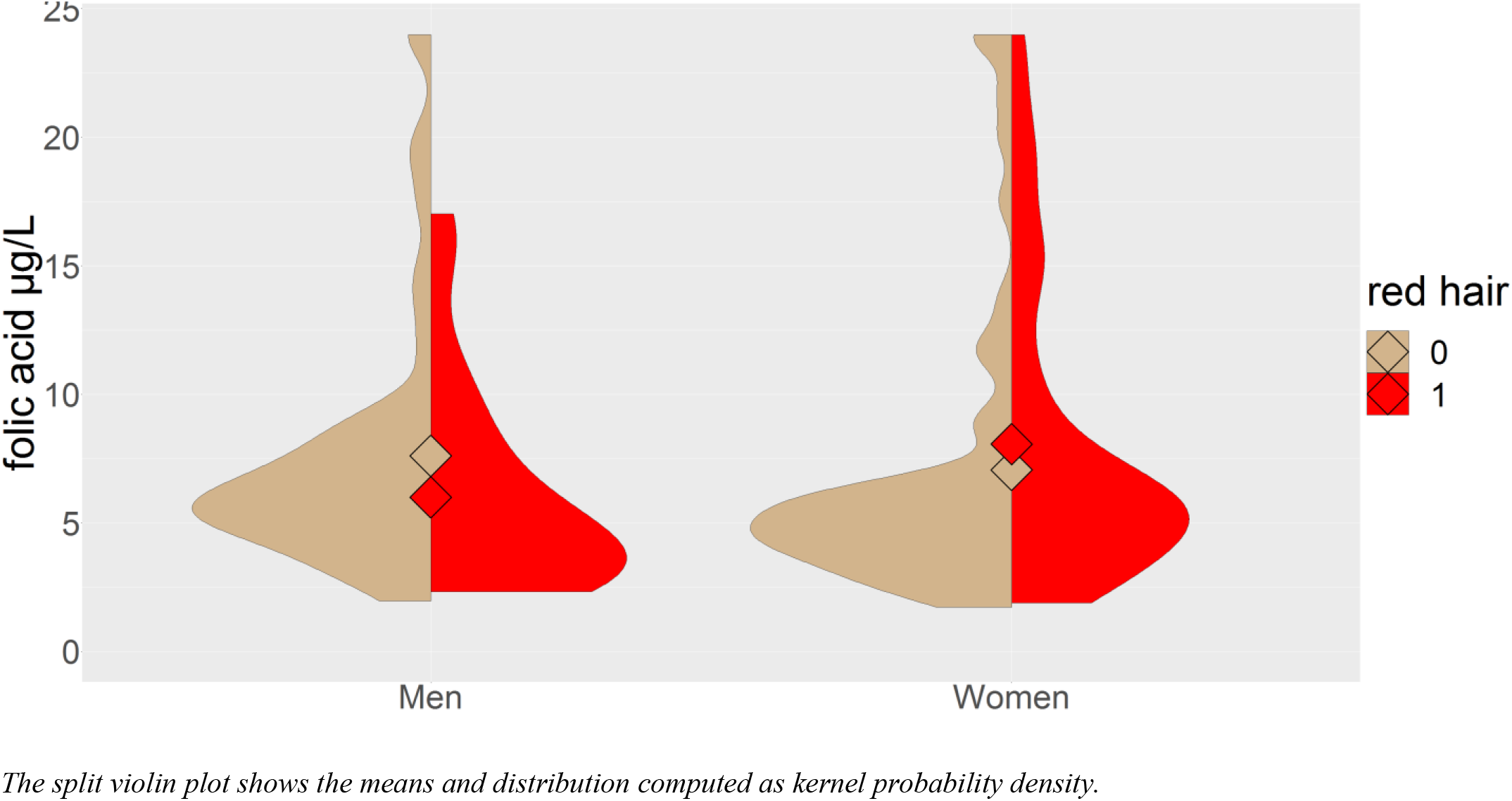
The effect of sex and red hair color on folic acid concentrations.

Separate partial Kendall analyses for redhaired and non-redhaired subjects showed that sun exposure had a minimal effect on calcidiol and folic acid concentrations in redhaired subjects (except for a negative effect of frequency of tanning salon visits on calcidiol levels). In non-redhaired subjects, sun exposure did have the expected effect on calcidiol and folic acid levels (see Figures 3–4, Table 3).

**Fig. 3.**
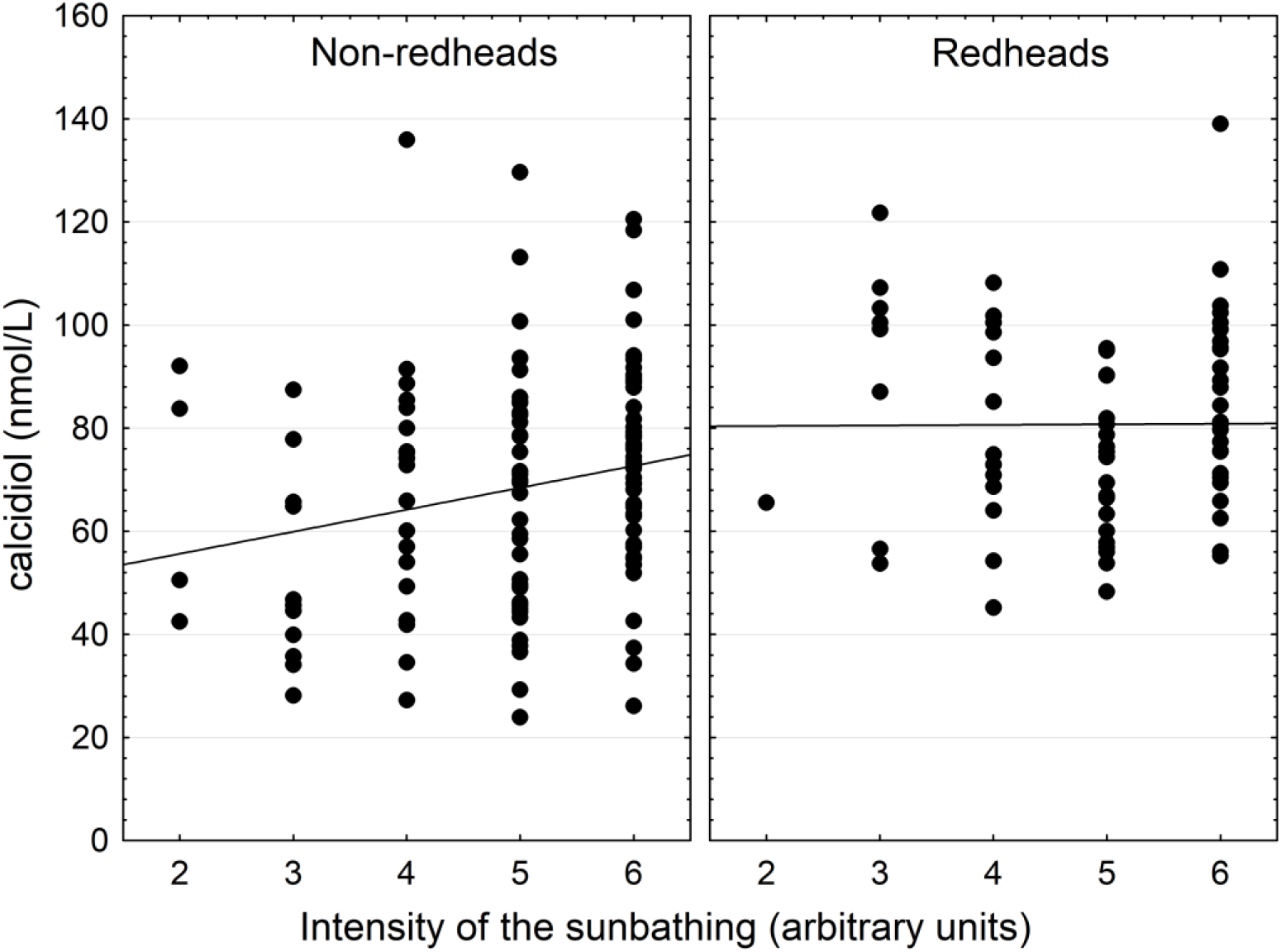
The effect of sun exposure on calcidiol concentrations.

**Fig. 4.**
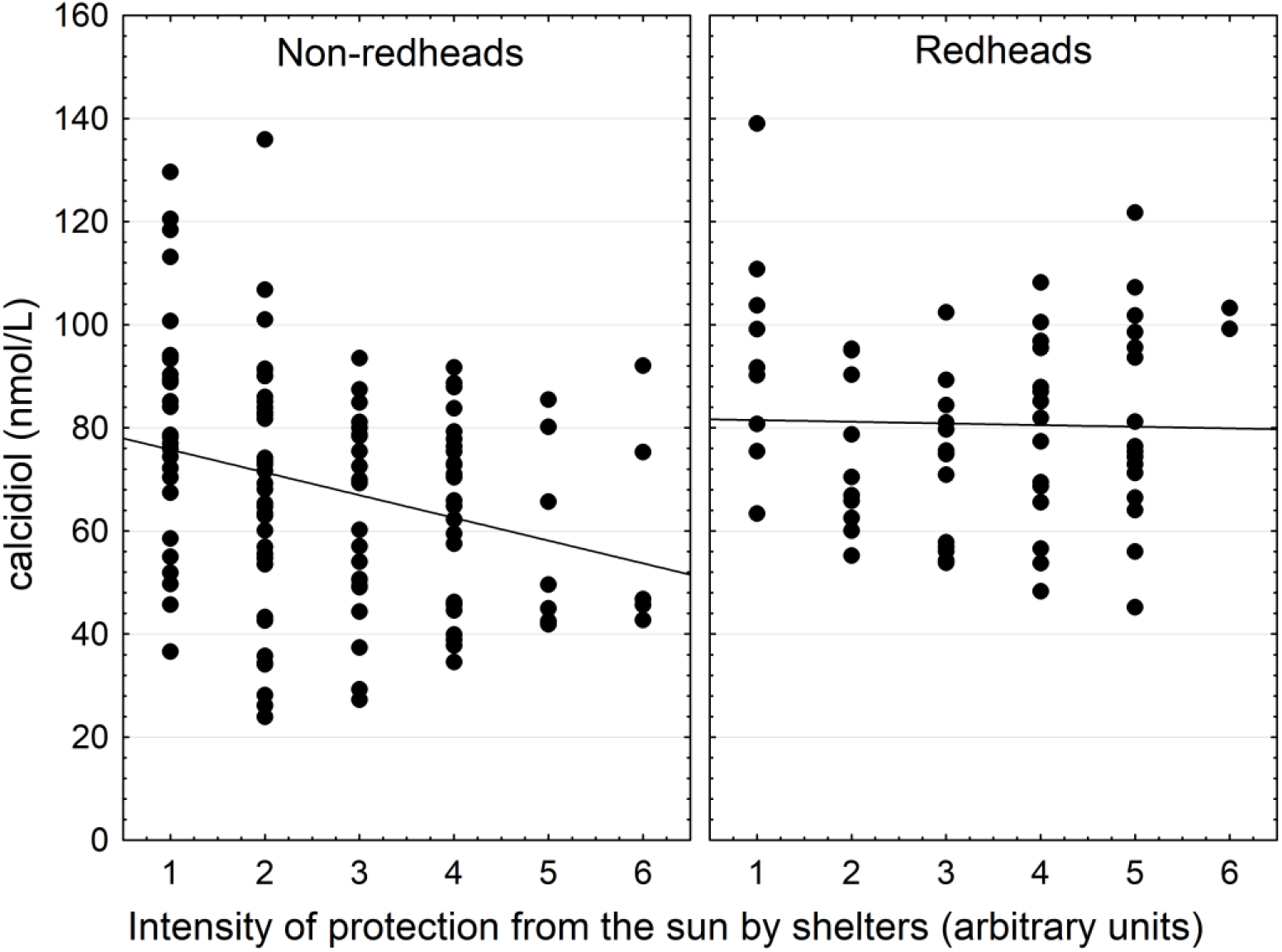
The effect of protection from sun exposure by seeking shelter on calcidiol concentrations.

**Table 3.** The effect of sun exposure-related variables on calcidiol and folic acid concentrations and suntan intensity.

|  | Redhaired subjects |  |  | Non-redhaired subjects |  |  |
| --- | --- | --- | --- | --- | --- | --- |
|  | calcidiol | folic acid | suntan | calcidiol | folic acid | suntan |
| Natural skin darkness | 0.015 | 0.097 | **0.475** | 0.091 | 0.050 | **0.431** |
| Suntan | 0.079 | -0.010 | NA | **0.241** | 0.098 | NA |
| Protection by sun creams | -0.073 | -0.061 | **-0.295** | 0.012 | -0.022 | **-0.145** |
| Protection by shelters | 0.050 | 0.068 | -0.156 | **-0.202** | -0.091 | **-0.296** |
| Sun exposure | 0.014 | 0.082 | **0.337** | **0.168** | -0.009 | **0.404** |
| Tanning salons | **-0.211** | -0.120 | 0.113 | 0.099 | **-0.161** | -0.079 |

*The table shows the strength and direction (Taus) of partial Kendall correlations (controlled for age and urbanization) between variables listed in the first column (see Material and methods) and calcidiol and folic acid concentrations as well as suntan. Significant correlations are in bold.*

## Discussion

Redhaired subjects had higher calcidiol concentrations and approximately the same folic acid concentrations as non-redhaired subjects. Results of partial correlations suggest that redhaired subjects need less sun exposure to achieve satisfactory calcidiol levels – and thereby probably also satisfactory levels of a biologically active vitamin D – than non-redhaired subjects do.

Differences between redhaired and non-redhaired subjects are likely to be due to differences in their physiology than an effect of their sunbathing-related behavior. For example, we observed no differences in the intensity of sun exposure between redhaired and non-redhaired subjects but redhaired subjects were less tanned at the time of the study. Redhaired subjects also reported that they use more intensive chemical and mechanical sun protection than their non-redhaired peers. In contrast to the situation in non-redhaired persons, redhaired persons’ calcidiol concentrations seemed independent of the intensity of sun exposure or protection from solar radiation. Redhaired subjects used vitamin D supplements less frequently but it should be noted that while the effect of these supplements on calcidiol levels was positive, it was at best modest and always non-significant. This absence of effect of vitamin D supplement use could be due to the fact that they tend to be used by persons with a diagnosed vitamin D deficiency.

Darker hues of natural hair but not of natural skin, both self-rated and measured spectrophotometrically, had a relatively strong negative effect on calcidiol concentrations. The two questions concerning natural skin hue and current tan were placed alongside each other in the questionnaire: it is therefore likely that responders rated the intensity of natural skin fairness or darkness as it looks untanned. The question on the darkness of natural hair, on the other hand, was in a different part of the questionnaire. It can be speculated that hair darkness actually reflects both the amount of eumelanin (positively) and intensity of sun exposure in the past (negatively). Calcidiol levels, meanwhile, could be negatively affected both by high eumelanin levels and by absence of sun exposure. With respect to skin (but not hair), sun exposure promotes darker hues. The opposite effect of eumelanin levels, which are positively correlated with darker natural skin hues and suntan intensity (acquired skin darkness), on calcidiol concentrations cancel each other. The result is an absence of correlation between darker skin hues and calcidiol levels.

It is known that solar radiation destroys folic acid by photolysis (Branda & Eaton, 1978; Jablonski & Chaplin, 2000). One could thus expect that folic acid concentrations would negatively correlate with the intensity of sun exposure and intensity of suntan. Actual data, however, show only weak positive correlations, none of which reach the formal level of significance. The only significant (and relatively strong) negative correlation with folic acid concentrations was found with respect to the frequency of visiting tanning salons. This pattern is in agreement with current theories (Jones, Lucock, Veysey, & Beckett, 2018) according to which in human populations there exist two mutually independent skin darkness latitudinal gradients, the results of two distinct selection pressures. The first gradient is found in populations which originated between subtropical and subpolar latitudes, that is, in the temperate climate. This gradient is the result of insufficient photosynthesis of vitamin D precursor in areas with low solar UV radiation. The second gradient is found in populations which originated between the tropics and the subtropics and its development was driven by excessive photolysis of folic acid in areas with intense solar radiation. The Czech Republic lies for the most part between 48° and 51° of northern latitude, where insufficient UV radiation rather than excess radiation could pose a problem, especially during the winter and spring months. It is indicative and perhaps clinically relevant that in our study, folic acid concentrations negatively correlated with the frequency of tanning salon visits.

We also found a rather strong positive correlation between the intensity of baldness and folic acid concentrations in men. Baldness intensity was not self-rated by women because our previous studies showed a minimal variability in this variable in young women. In men, however, both folic acid concentrations and baldness intensity strongly correlated with age. The strength of the correlation between folic acid concentration and baldness, however, was similar in cases where the age was (Tau = 0.21) and was not (Tau = 0.22) controlled for. In contrast to a general expectation, published data show no empirical evidence for an involvement of folic acid deficiency in alopecia (Almohanna, Ahmed, Tsatalis, & Tosti, 2019; Guo & Katta, 2017). Some studies even seem to support the notion of a positive association between folic acid and alopecia. For example, Rushton (2002) shows that among 200 healthy women complaining of increased hair shedding for over six months, only 1 had a “bellow range” folic acid level, while 57 had “above range” folic acid levels. Another study reported no significant difference in folate concentrations in a population of 91 female patients diagnosed with diffuse hair loss and 74 controls (Durusoy et al., 2009). Authors of that study did not, however, report folate concentrations in both groups, which may indicate that they had some unexpected results, such as lower folate concentrations in their controls.

As far as we know, our study is the first to have compared several methods of measuring the intensity of hair redness. Our results suggest that even the simplest method, i.e., the self-rating by participants, is satisfactory. Both methods of spectrophotometric measurement of hair redness worked similarly well, although the correlation between hair redness as estimated by subjects or other observers and hair redness as measured by the CIE L*a*b* color space method as the a* parameter (and possibly also the reflectance at 650 nm) was slightly higher than the correlation with redness as R calculated from reflectance according to Reed’s function. For example, partial Kendall correlation of self-rated hair redness with hair redness as a*, R, and reflectance at 650 nm was 0.528, 0.461, and 0.484, respectively. Similarly, correlation with calcidiol concentration was stronger for hair redness measured as the a* parameter than with hair redness measured as R (Table 2).

The main limitation of the present study is that our subjects cannot be considered a random sample of general Czech population. About half of the subjects who were asked to come to our laboratory to participate in an about 40-minute-long experiment politely refused. A few also refused to provide a blood sample for serological analysis. It is possible that persons who consented to participation and actually came to the experimental session form a specific population, for instance a group of highly altruistic subjects in good mood and good physical and mental condition. It is known that certain genetic and environmental factors influence variance more than physiological variables do (Flegr, 2013). Such factors may have, for example, negative effects on the health of a specific part of the population and positive effects on the health of others in the same population. If subjects who enjoy good health are more likely to be enrolled in the study (as may have been the case here), we may end up concluding that a particular factor, for instance redheadedness, has a positive effect on health and health-related variables although it has either no effect or even a negative effect on most members of a fully general population. Similarly, if subjects in poor health are more likely to be enrolled in a study – which is often the case with studies performed on patients with and without a particular disorder – a study can show that a particular factor has a negative effect on health although in majority of general population, its effect is positive. Our data suggest that such a sieve effect operated in our study, too. Firstly, latent infection with the common *Toxoplasma* parasite has a wide range of negative effects on the health of most members of the general population (Flegr & Escudero, 2016; Flegr, Prandota, Sovickova, & Israili, 2014). In our study, however, *Toxoplasma*-infected subjects enjoyed significantly better health than those who were *Toxoplasma*-free. (The effect of toxoplasmosis on health and wellbeing was a subject of another study performed on the same population of volunteers.) Secondly, a visual inspection of the violin plots for calcidiol and folic acid concentrations suggests that the distribution is truncated at the bottom and a subpopulation of individuals with a low concentration of these vitamins is missing from our sample. In a democratic country where people can refuse to participate in a study, the issue of non-representativeness of a sample due to sieve effect linked to the requirement of obtaining informed consent is hard or even impossible to avoid. It can be merely mitigated by making participation as easy and convenient as possible. It would be therefore most advisable to repeat our study on different populations of subjects who would not be selected or self-selected for better health.

## Conclusions

Based on previous observations of impaired health in fair-skinned people (Flegr & Sykorova, 2019), we predicted that redhaired subjects, who can be expected avoid sun exposure because of their sensitive skin, would have lower calcidiol levels. We confirmed that they indeed protect their skin from the sun by chemical and mechanical means. Nevertheless, we also found that in our self-selected sample, redhaired individuals did not avoid sun exposure any more than their non-redhaired peers and moreover, redhaired persons in our study had significantly higher calcidiol levels regardless of intensity of sun exposure. This discovery suggests that hair redness, the result of eumelanin synthesis downregulation, could be an evolutionary adaptation to life in higher latitudes where the photosynthesis of vitamin D precursor in skin is inadequate for large part of the year due to a low intensity of solar radiation. Our results suggest that redhaired individuals are capable of synthesizing sufficient amounts of calcidiol even when their sun exposure is minimal. Nonetheless, we should be cautious about generalizing this observation. This phenomenon was observed in two medium-sized samples of 93 men and 110 women who passed a relatively stringent self-selection process. Until this phenomenon is demonstrated in other, more representative populations, our conclusions must be considered merely preliminary.

## Acknowledgment

We would like to thank Anna Pilátová, Ph.D. for final revisions of our text. The study was supported by the Grant Agency of Charles University (project no. 1494218) and the Czech Science Foundation (grant No. 18-13692S).

## Data Accessibility

The final raw data set is available at figshare: https://figshare.com/s/50f5d6145b93a9892801

## Author contributions

JF, and KS designed research; KS, VF, JH, MB, LM, ŠK performed research, JF analyzed data and wrote the paper.

## Notes

https://figshare.com/s/50f5d6145b93a9892801

